# The relationship between subjective sleep quality and cognitive performance in healthy young adults: Evidence from three empirical studies

**DOI:** 10.1101/328369

**Authors:** Zsófia Zavecz, Nagy Tamás, Adrienn Galkó, Dezso Nemeth, Karolina Janacsek

## Abstract

The role of sleep in cognitive performance has gained increasing attention in neuroscience and sleep research in the recent decades, however, the relationship between subjective (self-reported) sleep quality and cognitive performance has not yet been comprehensively characterized. In this paper, our aim was to test the relationship between subjective sleep quality and a wide range of cognitive functions in a healthy young adult sample combined across three studies. Sleep quality was assessed by Pittsburgh Sleep Quality Index, Athens Insomnia Scale, and a sleep diary to capture general subjective sleep quality, and Groningen Sleep Quality Scale to capture prior night’s sleep quality. Within cognitive functions, we tested working memory, executive functions, and several sub-processes of procedural learning. To provide more reliable results, we included robust frequentist and Bayesian statistical analyses as well. Unequivocally across all analyses, we showed that there is no association between subjective sleep quality and cognitive performance in the domain of working memory, executive functions and procedural learning in healthy young adults. Our paper can contribute to a deeper understanding of subjective sleep quality and its measures, and we discuss various factors that may affect whether associations can be observed between subjective sleep quality and cognitive performance.

## Introduction

There is a widely accepted belief that experiencing lower sleep quality, including subjective experiences (e.g., reporting difficulties falling asleep, waking up frequently during the night, or feeling tired during the day)indisputably decreases cognitive performance. We can often hear people complaining about weaker memory and/or attentional performance in relation to their experienced sleep insufficiency. This phenomenon can be particularly prevalent amongst university students, where the pressure for academic performance is exceptionally high. The possible overestimation of the importance of one’s subjective sleep quality can lead to nocebo effects on cognitive performance. However, scientific evidence on the relationship between experienced subjective sleep quality and cognition is still lacking. Therefore our aim in the current study was to fill this gap and test whether subjective sleep quality is associated with cognitive performance in healthy young adults.

The role of sleep in cognitive performance has gained increasing attention in neuroscience and sleep research in the recent decades^1,2^. Numerous experimental methods exist that can be employed for examining the association between sleep and cognitive performance. Sleep parameters can be evaluated based on actigraph or electroencephalograph measurements (i.e., objective measures), which are time-consuming and require hardly accessible equipment. Hence researchers and clinicians still often tend to rely on questionnaires (i.e., subjective measures) to assess sleep parameters. This inclination has also motivated the current study to explore the relationship between sleep questionnaires and cognitive functions. It is important to note, that the relationship between objective sleep parameters and cognitive performance has been studied extensively, while the associations between subjective sleep quality and cognition have been largely neglected.

Previous studies have shown that subjective and objective sleep parameters could differ^3–5^. Subjective sleep quality can vary from the objective sleep quality because it is estimated by a combination of parameters, including the initiation of sleep, sleep continuity (number of awakenings), and/or depth of sleep. For instance, extreme deviations can occur between objective and subjective measures in case of sleep disorders, such as insomnia, or sleep-state misperception. According to Zhang and Zhao^6^, in sleep disorders, the objective and the subjective measures together should determine the type of treatment and medication. Stepanski, et al.^7^ showed that, within insomniac patients, the decisive factor whether a patient seeks medication is their subjective evaluation of their sleep quality and daytime functioning. Furthermore, in a placebo sleep study, Draganich and Erdal^8^ showed that assigned sleep quality predicted young adults’ performance in attentional and executive function tasks. Namely, participants were randomly told they had below average or above average sleep quality based on their brainwaves and other psychophysiological measures, and their belief about their sleep quality affected their cognitive performance. Thus, alongside therapeutic importance, the subjective evaluation of sleep quality could deepen our understanding of the complex relationship between sleep and cognitive performance. The aim of the present paper is to clarify the relationship between subjective sleep quality and aspects of cognitive functioning in healthy young adults.

One of the most widely-used sleep questionnaires is the Pittsburgh Sleep Quality Index (PSQI^9^), a self-administered questionnaire, in which participants rate their subjective sleep quality based on several questions, including the average amount of sleep during the night, the difficulty falling asleep, and other sleeping disturbances. Nevertheless, there are other popular measurements, such as the Athens Insomnia Scale (AIS^10^), which measures difficulties in falling asleep or maintaining sleep, and sleep diaries, which capture the sleeping habits of the participants from day to day, spanning a few days or weeks. Sleep questionnaires and sleep diaries are two different types of self-report measures: while a sleep questionnaire is retrospective, administered at a single point in time, and asks about various aspects of the sleep experience “in general”, sleep diary is an ongoing, daily self-monitoring. Libman, et al.^11^ showed that even though results of questionnaires and diaries are highly correlating, there are differences in the means of the sleep parameters depending on the type of self-report measurement. This suggests that the two measurement types are tapping the same domains but lead to somewhat different results due to methodological differences: questionnaires can be susceptible to memory distortion while sleep diaries may be distorted by atypical sleep experiences during the monitored period.

Previous research on subjective sleep quality and cognitive performance has led to mixed findings. While some studies focusing on healthy participants have shown that poorer sleep quality (measured by the PSQI score) was associated with weaker working memory performance^12^, executive functions^13^, and decision-making^14^, others have failed to find association between subjective sleep quality and cognitive performance ^7,15^. Focusing on sleep disorders, for instance, Naismith, et al.^16^ showed that greater subjective sleepiness was associated with weaker executive functions but not with IQ scores in patients with Obstructive Sleep Apnea. Importantly subjective sleepiness in this population was independent of polysomnographic sleep measures, which again suggests that even in sleep disorders subjective sleep quality may be an independent factor that underpins some aspects of cognitive functioning. Bastien, et al.^17^ showed different associations between subjective sleep quality and cognitive performance in patients with insomnia with and without treatment and in elderly participants who reported good sleep quality. Interestingly, in good sleepers, greater subjective depth, quality, and efficiency of sleep was associated with better performance on attention and concentration tasks but *poorer* memory performance, calling for further studies to test the complex relationship between subjective sleep quality and aspects of cognitive functioning.

Importantly, these previous studies focused on diverse populations, including adolescents, elderly and clinical groups, and relied on sample sizes ranging from around 20 to 100, with smaller sample sizes potentially limiting the robustness of the observed results. A recent powerful meta-analysis focused on one aspect of sleep quality, namely, on the effect of self-reported sleep duration on cognitive performance in elderly participants, and reported that both short and long sleep increased the odds of poor cognitive performance^18^. Systematic investigations on the relationship between other aspects of subjective sleep quality and cognitive performance using larger sample sizes are, however, still lacking. Here we aimed to, at least partly, fill this gap by testing the associations between subjective sleep quality and cognitive performance using a more detailed examination of subjective sleep quality and a wide range of neuropsychological tests in a relatively large sample of healthy young adults.

In previous investigations focusing on the association between subjective sleep quality and various aspects of cognitive performance, the potential relationship with procedural learning/memory has largely been neglected. The procedural memory system underlies the learning, storage, and use of cognitive and perceptual-motor skills and habits^19^. Evidence suggests that the system is multifaceted in that it supports numerous functions that are performed automatically, including sequences, probabilistic categorization, and grammar, and perhaps aspects of social skills^20–24^. In light of the importance of this memory system, the clarification of its relationship with sleep would be indispensable.

In this paper, our aim was to provide an extensive investigation on the relationship between subjective sleep quality and cognitive performance, including procedural learning in healthy young adults. To increase the robustness of our analyses, we created a database of 235 participants’ data by combining three separate datasets from our lab. Subjective sleep quality was assessed not only by PSQI but also by other, less frequently used sleep quality measures: namely, Athens Insomnia Scale (AIS, Study 1-3), Groningen Sleep Quality Scale (GSQS, Study 2), and a sleep diary (Study 2). These separate measures capture somewhat different aspects of self-reported sleep quality and thus provide a detailed picture. In all three studies working memory, executive functions and several sub-processes of procedural learning were probed. This approach enabled us to test – within the same participants and same experimental designs– whether procedural learning is differentially associated with subjective sleep quality as opposed to working memory and executive functions. To test the amount of evidence either for associations or no associations between subjective sleep and cognitive performance in the study population, we calculated Bayes Factors that offers a way of evaluating evidence against or in favor of the null hypothesis, respectively. To the best of our knowledge, this is the first extensive investigation on the relationship between subjective sleep quality and cognitive functions, covering such a wide range of assessments, in healthy young adults.

## Methods

### Participants

Participants were selected from a large pool of undergraduate students from Eötvös Loránd University in Budapest. The selection procedure was based on the completion of an online questionnaire assessing mental and physical health status. Respondents reporting current or prior chronic somatic, psychiatric or neurological disorders, or the regular consumption of drugs other than contraceptives were excluded. In addition, individuals reporting the occurrence of any kind of extreme life event (e.g., accident) during the last three months that might have had an impact on their mood, affect and daily rhythms were not included in the study.

The data was obtained from three different studies with slightly different focus. Importantly, the analyses presented in the current paper are completely novel, none of the separate studies focused on the relationship between subjective sleep quality and cognitive performance. Forty-seven participants took part in Study 1^25^, 103participants took part in Study 2^26^, and 85 participants took part in Study 3^27^. Descriptive characteristics of participants in the three studies are listed in Table 1. All participants provided written informed consent and received course credits for taking part in one of the studies. The studies were approved by the Research Ethics Committee of Eötvös Loránd University, Budapest, Hungary (201410, 2016/209). The study was conducted in accordance with the Declaration of Helsinki.

**Table 1.** Descriptive characteristics of participants.

| **Study** | **n** | **Age**<br>***M (SD)*** | **Sex** | **Years in education**<br>***M (SD)*** |
| --- | --- | --- | --- | --- |
| Study 1 | 47 | 21.38 (1.79) | 10M/37F | 14.36 (1.58) |
| Study 2 | 103 | 21.62 (2.00) | 30M/73F | 14.50 (1.74) |
| Study 3 | 85 | 20.99 (1.59) | 23M/62F | 14.28 (1.60) |

*Note* M - male, F - female

### Procedure

Three separate studies on the association of subjective sleep quality (assessed by sleep questionnaires) and procedural learning (measured with ASRT) and other cognitive functions, such as working memory and executive functions were conducted. The tasks and questionnaires included in the studies and the timing of the ASRT task slightly differed. In Study 2, we included additional subjective sleep questionnaires: (1) a sleep diary to assess day-to-day general sleep quality and (2) Groningen Sleep Quality Scale (GSQS) to assess prior night’s sleep quality.

In all three studies, PSQI and AIS sleep quality questionnaires were administered online, while the GSQS in Study 2 and the tasks assessing cognitive performance in all studies were administered in a single session in the lab. To ensure that participants do the tests in their preferred time of the day, the timing of the session was chosen by the participants themselves (between 7 am and 7 pm). The timing of the sessions was normally distributed in all three studies, with most participants performing the tasks during daytime between 11 am and 3 pm. The sleep diary in Study 2 was given to the participants 1 to 2 weeks prior to the cognitive assessment.

### Questionnaires and tasks

All cognitive performance tasks and subjective sleep questionnaires are well-known and widely used in the field of psychology and neuroscience (for details about each task and questionnaire, see Supplementary methods).

#### Subjective sleep quality questionnaires

– To capture general sleep quality, we administered the *Athens Insomnia Scale (*Hungarian version: ^28^), the *Pittsburgh Sleep Quality Index (* Hungarian version: ^29^), and a *Sleep diary* ^30^. Additionally, to capture the sleep quality of the night prior testing, we administered the *Groningen Sleep Quality Scale (*GSQS)^31^, (Hungarian version: ^32^).

#### Cognitive performance tasks

– *Working memory* was measured by the Counting Span task^33–35^ (Hungarian version: ^36^) and *executive functions* were assessed by the Wisconsin Card Sorting Test (WCST) ^37,38^, on a Hungarian sample: ^39^. The outcome measure of this task was the number of *perseverative errors*, which shows the inability/difficulty to change the behavior despite feedback. *Procedural learning* was measured by the explicit version of the Alternating Serial Reaction Time (ASRT) task (FigureS1, see also ^40^). There are several learning indices that can be acquired from this task. *Higher-order sequence learning* refers to the acquisition of the sequence order of the stimuli. *Statistical learning* refers to the acquisition of frequency information embedded in the task. However, previous ASRT studies often assessed *Triplet learning*, which is a mixed measure of acquiring frequency and sequential information (for details, see Supplementary methods). In addition to these learning indices, we measured the average reaction times (RTs) and accuracy (ACC), and changes in RT and ACC performance from the beginning to the end of the task, that indicate g*eneral skill learning*, such as more efficient visuo-motor and motor-motor coordination as the task progresses^41^.

### Data analysis

Statistical analyses were conducted in R 3.5.2^42^ with the lme4^43^ and robustlmm^44^ packages.

#### Analysis of the ASRT data

Performance in the ASRT task was analyzed by repeated measures analyses of variance (ANOVA) in each study (for details of these analyses, see Supplementary methods). Based on these ANOVAs, *Triplet learning, Higher-order sequence learning* and *Statistical learning* occurred in all three studies, both in ACC and RT (all *p*s <.001, for details, see Supplementary results, and Figure S2).

#### Analysis of the relationship between subjective sleep quality and cognitive performance

Subjective sleep quality scales (PSQI and AIS) were combined into a single metric, using principal component analysis. Then separate linear mixed-effect models were created for each outcome measure (i.e., performance metric), where the aggregated sleep quality metric (hereinafter referred to as sleep disturbance) was used as a predictor, and ‘Study’ (1, 2 or 3) was added as random intercept. This way we could estimate an aggregated effect while accounting for the potential differences between studies. As residuals did not show normal distribution, we used robust estimation of model parameters using the robustlmm package^44^. Bayes Factors (BF_01_) were calculated by using the exponential of the Bayesian Information Criterion (BIC) of the fitted models minus the BIC of the null models – that contained no predictor, but a random intercept by study^45^. The BF is a statistical technique that helps conclude whether the collected data favors the null-hypothesis (*H*_0_) or the alternative hypothesis (*H*_1_); thus, the BF could be considered as a weight of evidence provided by the data^46^. It is an effective mathematical approach to show if there’s no association between two measures. In Bayesian correlation analyses, *H*_0_ is the lack of associations between the two measures, and *H*_1_ states that association exists between the two measures. Here we report BF_01_ values. According to Wagenmakers et al.^46^, BF_01_ values between 1 and 3 indicate anecdotal evidence for *H*_0_, while values between 3 and 10 indicate substantial evidence for *H*_0_. Conversely, while values between 1/3 and 1 indicate anecdotal evidence for *H*_1_, values between 1/10 and 1/3 indicate substantial evidence for *H*_1_. If the BF is below 1/10, 1/30, or 1/100, it indicates strong, very strong, or extreme evidence for *H*_1_, respectively. Values around one do not support either *H*_0_ or *H*_1_. Thus, Bayes Factor is a valuable tool to provide evidence for no associations between constructs as opposed to frequentists analyses, where no such evidence can be obtained based on non-significant results.

In Study 2, to test the association between the additional subjective sleep quality measures and cognitive performance, we used a similar robust linear regression, this time without random effects. Bayes factors were calculated in the previously described way. Normality of data distribution was violated in sleep questionnaire scores, thus we only used robust methods.

## Results

### Combining sleep quality metrics

Principal component analysis was used to combine PSQI and AIS into a single ‘sleep disturbance’ metric. The Bartlett’s test of sphericity indicated that the correlation between the scales was adequately large for a PCA, χ^2^(235) = 84.88, p < .0001. One principal factor with an eigenvalue of 1.55 was extracted to represent sleep disturbance. The component explained 83.7% of the variance, and it was named ‘sleep disturbance’, as higher values of this metric show more disturbed sleep.

### Associations between subjective sleep quality and cognitive performance

As described above, to study the associations between subjective sleep quality and cognitive performance, separate linear mixed-effect models were created for each outcome measure (i.e., cognitive performance metric), where sleep disturbance was used as a fixed predictor, and ‘Study’ was added as random intercept. Sleep disturbance did not show an association with any of the cognitive performance metrics (see Table 2 and Figure 1). Bayes Factors ranged from 3.49 to 14.65, thus, there is *substantial* evidence for *no* association between subjective sleep quality and the measured cognitive processes^46^.

**Table 2.** The association of sleep disturbance with cognitive performance metrics.

| Outcome | $\beta$ | Std.error | *t* | df | *p* | BF<sub>01</sub> |
| --- | --- | --- | --- | --- | --- | --- |
| **ACC learning indices** |  |  |  |  |  |  |
| ACC Triplet learning | -0.061 | 0.053 | -1.14 | 230 | 0.2552 | 5.09 |
| ACC Higher-order sequence learning | -0.072 | 0.055 | -1.31 | 230 | 0.1904 | 7.00 |
| ACC Statistical learning | -0.026 | 0.059 | -0.43 | 230 | 0.6657 | 13.12 |
| **RT learning indices** |  |  |  |  |  |  |
| RT Triplet learning | -0.006 | 0.036 | -0.17 | 230 | 0.8620 | 10.86 |
| RT Higher-order sequence learning | 0.011 | 0.028 | 0.38 | 230 | 0.7070 | 14.28 |
| RT Statistical learning | -0.036 | 0.064 | -0.55 | 230 | 0.5812 | 13.28 |
| **General skill indices** |  |  |  |  |  |  |
| Average ACC | 0.039 | 0.034 | 1.15 | 230 | 0.2529 | 3.49 |
| ACC general skill learning | 0.068 | 0.035 | 1.93 | 230 | 0.0543 | 3.68 |
| RT average | -0.066 | 0.066 | -1.00 | 230 | 0.3192 | 11.47 |
| RT general skill learning | -0.059 | 0.060 | -0.98 | 230 | 0.3278 | 6.61 |
| **WM and EF indices** |  |  |  |  |  |  |
| Counting Span | -0.047 | 0.064 | -0.73 | 230 | 0.4641 | 14.65 |
| WCST – perseverative error | 0.035 | 0.051 | 0.67 | 224 | 0.5007 | 8.92 |

*Note:* The table shows standardized regression coefficients for sleep disturbance, where the ‘Study’ random intercept was included in separate linear mixed-effect models for each cognitive performance metrics. BF<sub>01</sub> was
derived from BIC (see text for details). ACC = accuracy. RT = reaction time. WM = working memory. EF = executive function. WCST = Wisconsin Card Sorting Test.

**Figure 1.**
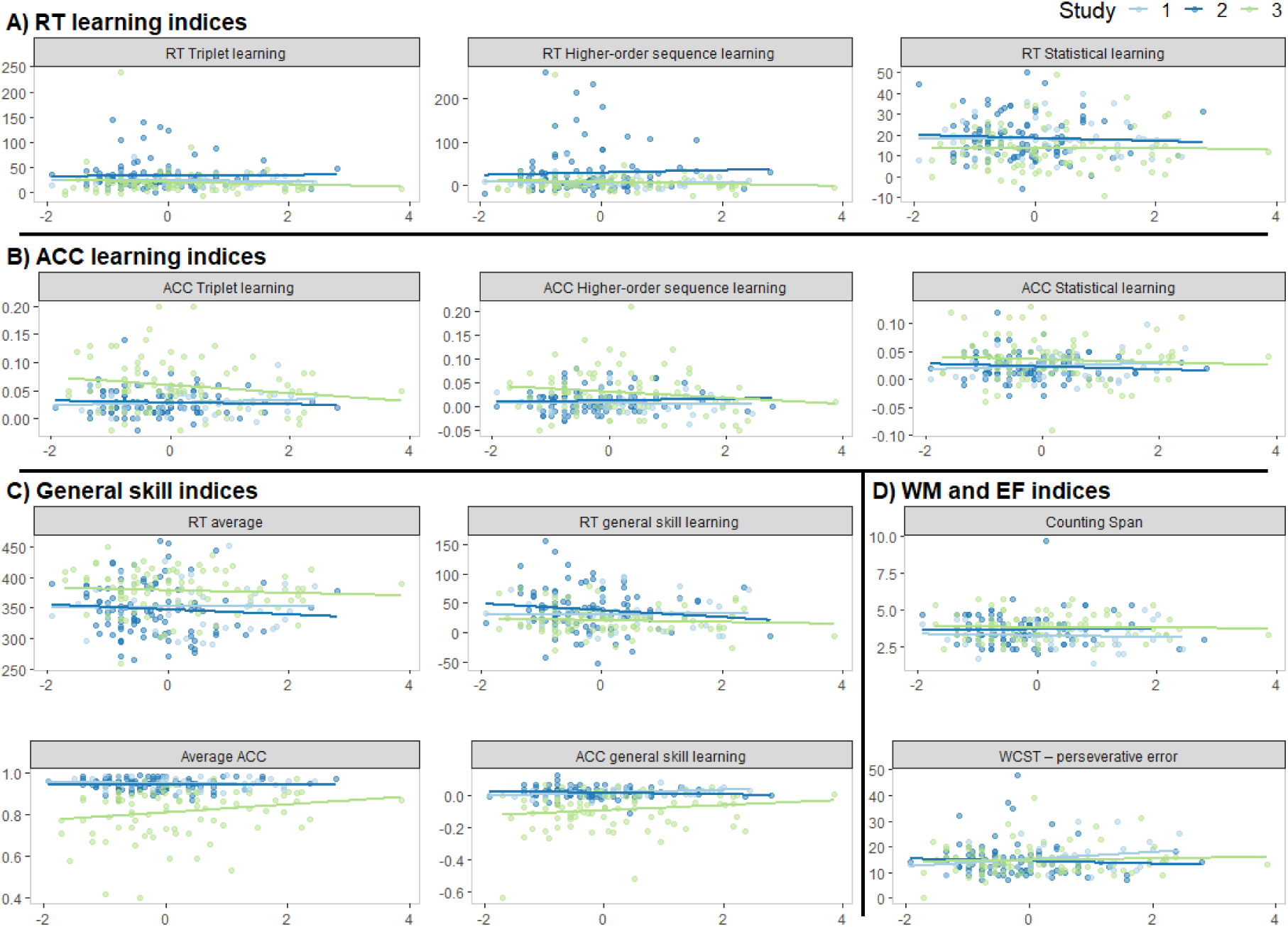
Association between sleep disturbance and cognitive performance metrics by study. The scatterplots and the linear regression trendlines show no association between subjective sleep quality and procedural learning indices in terms of reaction time (RT, A), or accuracy (ACC, B), general skill indices in terms of RT or ACC (C), and working memory and executive function indices (D).

In **Study 2**, to study the associations between further subjective sleep quality questionnaires and cognitive performance, we created a separate linear mixed-effect models for each outcome measure (i.e., cognitive performance metric), and each additional sleep questionnaire (e.g. sleep diary and GSQS). Sleep diary scores did not show association with any of the cognitive performance metrics (all *p*s > .10, see Table 3 and Figure 2). Bayes Factors ranged from 2.41 to 12.58, thus, there is evidence for *no* association between subjective sleep quality and the measured cognitive processes ^46^.

**Table 3.** The association of sleep diary with cognitive performance metrics.

| **Outcome** | **$\beta$** | **Std.error** | **$t$** | **df** | **$p$** | **BF<sub>01</sub>** |
| --- | --- | --- | --- | --- | --- | --- |
| **ACC learning indices** |  |  |  |  |  |  |
| ACC Triplet learning | -0.061 | 0.093 | -0.65 | 97 | 0.5159 | 7.00 |
| ACC Higher-order sequence learning | -0.065 | 0.091 | -0.72 | 97 | 0.4763 | 8.60 |
| ACC Statistical learning | 0.013 | 0.095 | 0.14 | 97 | 0.8916 | 10.40 |
| **RT learning indices** |  |  |  |  |  |  |
| RT Triplet learning | -0.044 | 0.060 | -0.73 | 97 | 0.4673 | 7.36 |
| RT Higher-order sequence learning | 0.005 | 0.049 | 0.10 | 97 | 0.9176 | 10.32 |
| RT Statistical learning | -0.158 | 0.096 | -1.65 | 97 | 0.1027 | 7.01 |
| **General skill indices** |  |  |  |  |  |  |
| Average ACC | 0.042 | 0.104 | 0.40 | 97 | 0.6879 | 8.70 |
| ACC general skill learning | -0.141 | 0.080 | -1.76 | 97 | 0.0823 | 2.41 |
| RT average | -0.105 | 0.103 | -1.02 | 97 | 0.3102 | 12.58 |
| RT general skill learning | -0.052 | 0.093 | -0.56 | 97 | 0.5777 | 7.63 |
| **WM and EF indices** |  |  |  |  |  |  |
| Counting Span | -0.053 | 0.089 | -0.59 | 97 | 0.5549 | 6.73 |
| WCST – perseverative error | -0.055 | 0.064 | -0.86 | 97 | 0.3911 | 6.62 |

*Note:* The table shows standardized regression coefficients for sleep diary scores in separate linear mixed-effect models for each cognitive performance metrics. BF<sub>01</sub> was derived from BIC (see text for details). ACC = accuracy. RT = reaction time. WM = working memory. EF = executive function. WCST = Wisconsin Card Sorting Test.

**Figure 2.**
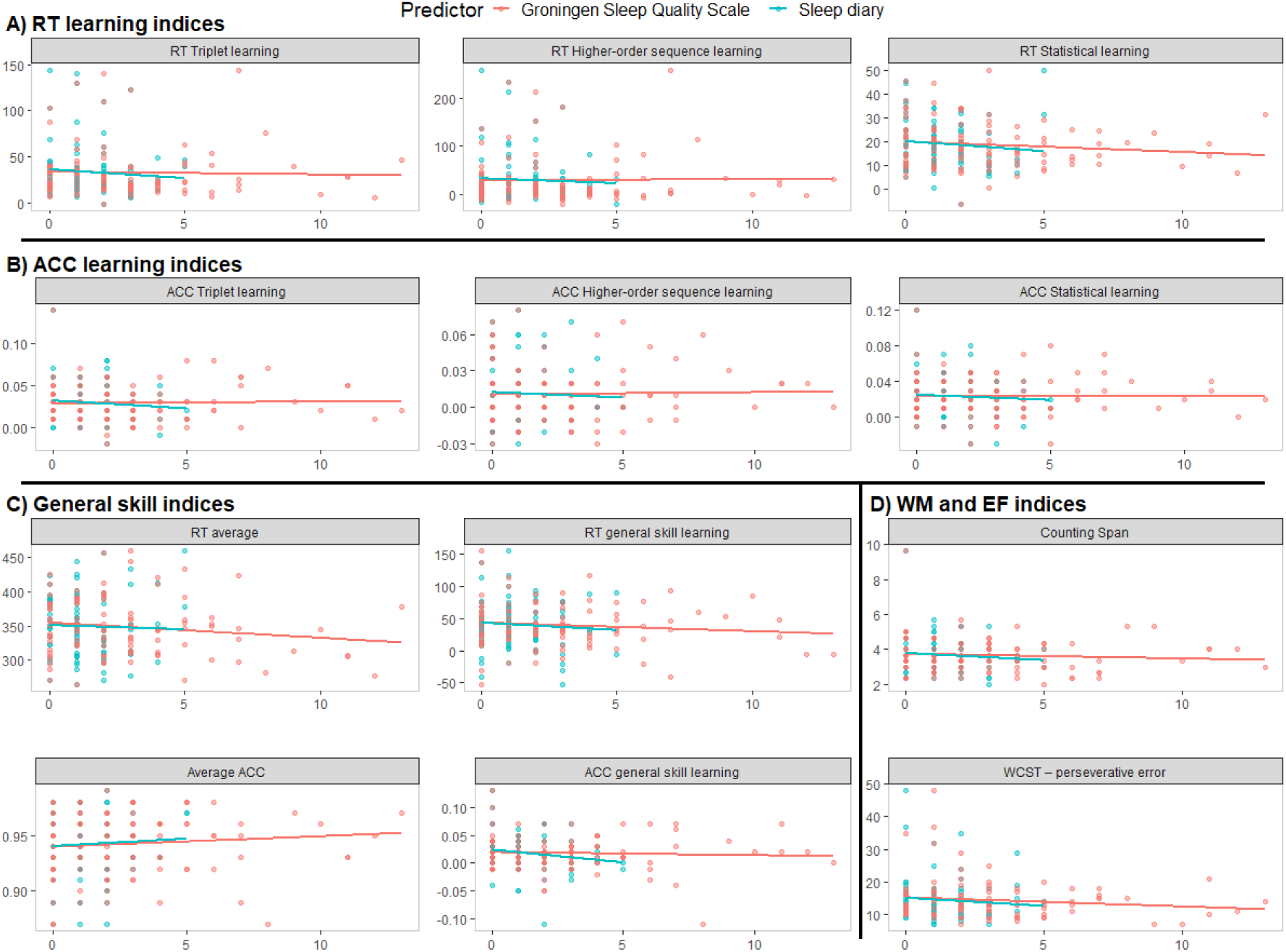
Association between sleep diary and GSQS scores and cognitive performance metrics. The scatterplots and the linear regression trendlines show no association between subjective sleep quality (measured with a sleep diary (blue) or the GSQS (red)) and procedural learning indices in terms of reaction time (RT, A), or accuracy (ACC, B), general skill indices in terms of RT or ACC (C), and working memory and executive function indices (D).

Similarly, GSQS scores did not show association with any of the cognitive performance metrics (all *p*s > .25, see Table 4 and Figure 2). Bayes Factors ranged from 3.30 to 11.85, thus, there is *substantial* evidence for *no* association between subjective sleep quality and the measured cognitive processes ^46^.

**Table 4.**
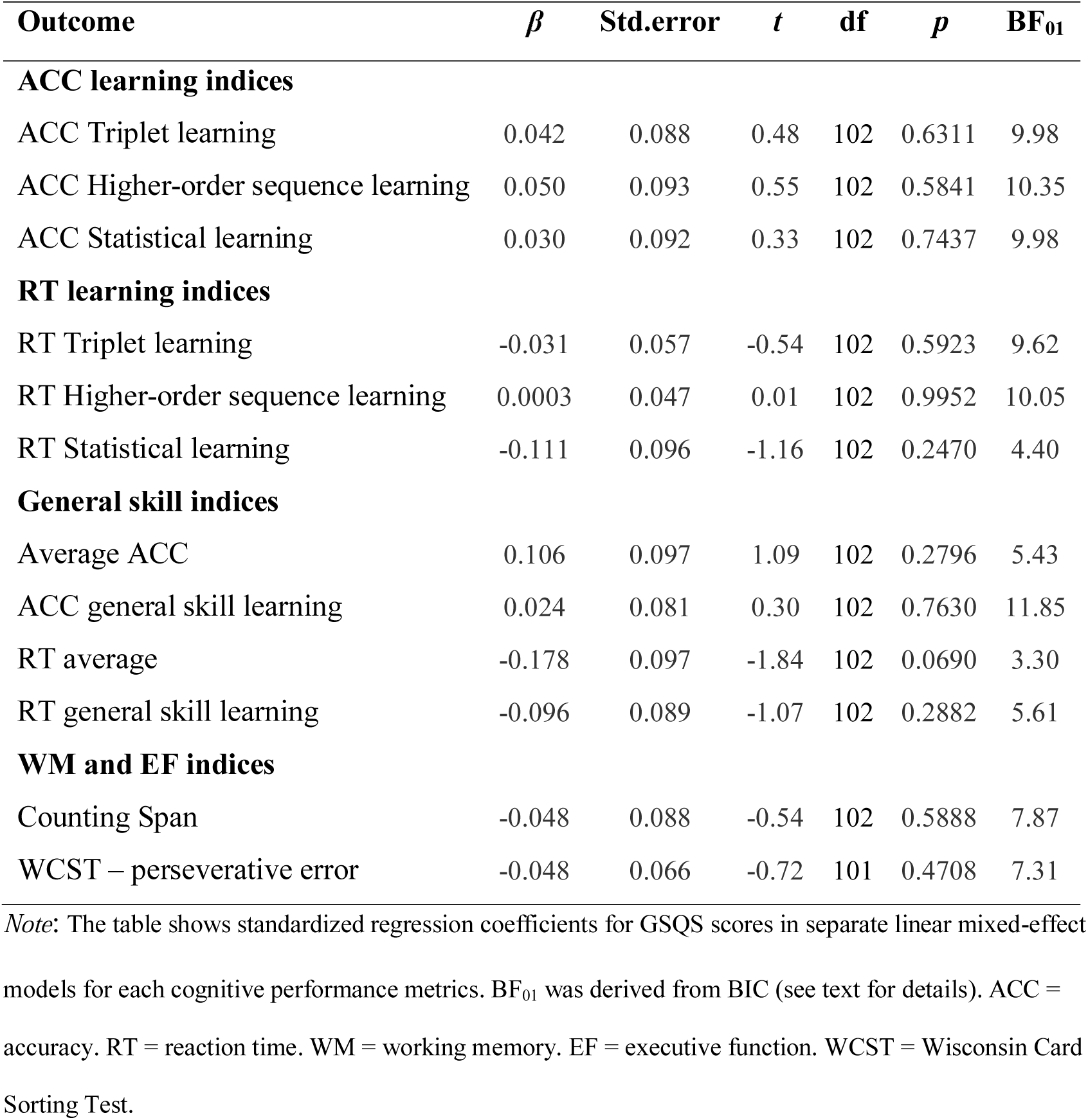
The association of GSQS with cognitive performance metrics.

| **Outcome** | **$\beta$** | **Std.error** | **$t$** | **df** | **$p$** | **BF<sub>01</sub>** |
| --- | --- | --- | --- | --- | --- | --- |
| **ACC learning indices** |  |  |  |  |  |  |
| ACC Triplet learning | 0.042 | 0.088 | 0.48 | 102 | 0.6311 | 9.98 |
| ACC Higher-order sequence learning | 0.050 | 0.093 | 0.55 | 102 | 0.5841 | 10.35 |
| ACC Statistical learning | 0.030 | 0.092 | 0.33 | 102 | 0.7437 | 9.98 |
| **RT learning indices** |  |  |  |  |  |  |
| RT Triplet learning | -0.031 | 0.057 | -0.54 | 102 | 0.5923 | 9.62 |
| RT Higher-order sequence learning | 0.0003 | 0.047 | 0.01 | 102 | 0.9952 | 10.05 |
| RT Statistical learning | -0.111 | 0.096 | -1.16 | 102 | 0.2470 | 4.40 |
| **General skill indices** |  |  |  |  |  |  |
| Average ACC | 0.106 | 0.097 | 1.09 | 102 | 0.2796 | 5.43 |
| ACC general skill learning | 0.024 | 0.081 | 0.30 | 102 | 0.7630 | 11.85 |
| RT average | -0.178 | 0.097 | -1.84 | 102 | 0.0690 | 3.30 |
| RT general skill learning | -0.096 | 0.089 | -1.07 | 102 | 0.2882 | 5.61 |
| **WM and EF indices** |  |  |  |  |  |  |
| Counting Span | -0.048 | 0.088 | -0.54 | 102 | 0.5888 | 7.87 |
| WCST – perseverative error | -0.048 | 0.066 | -0.72 | 101 | 0.4708 | 7.31 |

*Note:* The table shows standardized regression coefficients for GSQS scores in separate linear mixed-effect
models for each cognitive performance metrics. BF<sub>01</sub> was derived from BIC (see text for details). ACC = accuracy. RT = reaction time. WM = working memory. EF = executive function. WCST = Wisconsin Card Sorting Test.

## Discussion

Our aim was to investigate, in healthy young adults, the relationship between subjective sleep quality (assessed by self-report measures) and performance in various cognitive functions, such as working memory, executive functions, and procedural learning (which has mainly been neglected in studies of subjective sleep quality before). While the relationship between objective sleep parameters and cognitive performance has been widely studied, the associations between subjective sleep quality and cognition have been largely neglected. To provide more reliable results, we combined data obtained from three different studies and included robust frequentists and Bayesian statistical analysis as well. We did not find associations between subjective sleep quality and cognitive performance. Moreover, the Bayes factors provided substantial evidence for no associations between subjective sleep quality and most measures of procedural learning, or other cognitive performance measure included in our investigation.

None of the procedural learning indices showed associations with subjective sleep quality (supported by Bayes Factors). Higher-order sequence learning, Statistical learning, Triplet learning and general skill learning (both in terms of ACC and RT) thus seem to be independent of self-reported sleep quality. In procedural learning, its relationship with objective sleep quality is still debated ^47–50^. The results so far have been controversial, as some studies have shown associations between various aspects of sleep, such as time spent in rapid eye movement (REM) sleep^47^ or time spent in non-rapid eye movement (NREM) 2 sleep^48^ and procedural learning, while others, focusing primarily on patients with sleep disorders, or examining sleep effects in an AM-PM vs. PM-AM design have not found such associations ^49–53^. Here we focused on subjective sleep quality and showed evidence for no association with procedural learning in healthy young adults, which is consistent with previous studies showing no relationship between procedural learning performance and objective sleep measures^49–53^. Importantly, there is great variability across studies in using different tasks or testing different sleep parameters, as well as in other study settings (e.g., population characteristics, or time of the day of testing). Our results suggest that procedural learning is not related to subjective sleep quality in healthy young adults. Nevertheless, further investigations with similar settings (e.g., with the same tasks and/or same sleep parameters) across studies are needed to clarify the specific circumstances under which subjective and/or objective sleep quality may be associated with aspects of procedural learning.

Contrary to our expectations, working memory and executive functions also did not show association with subjective sleep quality. As presented in the introduction, some studies reported associations between subjective sleep quality and working memory performance^12^, executive functions^13^ and decision making^14^, although other studies also exist that failed to find such associations ^7,15^. These studies focused primarily on healthy/disordered elderly or adolescent populations. To the best of our knowledge, our study is the first that investigates cognitive performance in association with subjective sleep quality in a relatively large sample of healthy young adults. A possible explanation for the diverse results is that when cognitive performance peaks in young adulthood, subjective sleep quality may not have a substantial impact on it, while in other populations, such as in adolescents, older adults, or in various clinical disorders, where cognitive performance has not yet peaked or have declined, subjective sleep quality can have a bigger impact on performance. In line with this explanation, Saksvik, et al.^54^ found in their meta-analysis that young adults are not as prone to the negative consequences of shift work as the elderly.

It is also worth noting that sleep quality disturbance is more prevalent in adolescent or elderly populations and in clinical disorders. Consequently, variance and extremities in subjective sleep quality could be greater in these populations, while it can remain relatively low in healthy young adults. However, the variance of the cognitive performance tasks and the subjective sleep questionnaires scores in this paper were sufficient to test associations between subjective sleep quality and cognitive performance (for details, see Supplementary results). Thus, we believe that finding no association between subjective sleep quality and cognitive performance in the current study is not due to methodological issues (such as low variance of the used measures). Another possibility that may affect the relationship between subjective sleep quality and cognitive performance is whether a poorer sleep quality is relatively transient or persists for years or even for decades. It is plausible and would worth to test *systematically* whether, and in which cases, such long-term poor sleep quality has a more detrimental effect on cognitive performance compared to a relatively more recent decline in sleep quality.

Associations between objective sleep quality (measured by actigraph or electroencephalograph) and various aspects of memory or executive functions have been frequently reported before^1,2^. Here we showed that subjective sleep quality is not associated with working memory and executive functions. As already mentioned in the Introduction, this dissociation suggests that subjective and objective sleep quality, although measuring the same domains, do not necessarily measure the same aspects of sleep quality and sleep disturbances. Landry, et al.^3^ compared a sleep questionnaire (namely, PSQI) and a sleep diary with actigraphy data. According to their results, while the sleep questionnaire and the sleep diary scores moderately correlated, actigraphy data had only weak correlation with both self-reported measures. Guedes, et al.^4^ showed that the discrepancy of sleep duration quantified by actigraphy or self-reported measures can even be 1-2 hours on average. Objective and subjective assessments of sleep quality, despite the fact that they often carry labels that imply direct relationship or equivalence, may relate to different parameters^5^, such as impressions of sleep quality, restedness, or sleep depth do not appear to be strongly correlated with sleep architecture. Furthermore, subjective sleep quality might be represented by a combination of more than one objective sleep parameter.

It is also possible that the important parameters of sleep that contribute to memory or executive function performance cannot be captured with self-reported instruments. For example, it is often reported (see also above), that spindle activity or time spent in slow-wave sleep (SWS), or in REM sleep is essential for memory consolidation^55–57^. These sleep parameters could not be evaluated subjectively. Also, in laboratory sleep examinations, the general subjective sleep quality together with the sleep quality of preceding days of the examination is usually carefully controlled. Thus, potentially, the parameters showed to be important in the associations of objective sleep parameters and cognitive performance during a given night can only be measured in these carefully controlled conditions (i.e., when sleep quality in general and in preceding days are good). Hence it is possible that while results with objective sleep quality show how healthy sleep is related to cognitive functioning, results with subjective sleep quality may reflect aspects of sleep disturbances and their potential relationship with cognitive functioning.

Importantly, we found no associations with cognitive performance both for general sleep quality (assessed for a one-month period) and for the previous night’s subjective sleep quality. The Bayes Factors showed evidence for no associations between previous night’s sleep quality and procedural learning, working memory or executive function, in the case of the GSQS questionnaire. These results suggest that in healthy young adults neither persistent nor transient subjective sleep quality contribute to cognitive performance.

Considering the dissociation between objective and subjective sleep quality, the use of self-reported tools to measure sleep quality should be treated carefully in generalizing results to all aspects of sleep quality. Usage of these questionnaires should also be avoided for diagnostic purposes, as also suggested by West, et al.^58^, who attempted to validate sleep questionnaires with PSG in insomniac patients. Comparative studies have also shown significant discrepancies between subjective and objective measures of sleep pathology^6,59^. Subjective sleep quality, rather than used interchangeably with objective sleep quality, should be assessed to gain further information of participants’ or patients’ sleep, as it may have different explanatory and predictive value to cognitive performance, and treatment-seeking or outcome^7,8^. Subjective sleep quality can be especially informative in populations with extremities in subjective sleep experience that are more susceptible to sleep disturbance. For instance, Gavriloff, et al. ^60^ in a recent paper showed that providing sham feedback to patients with insomnia influenced their daytime symptoms and cognitive performance such as attention and vigilance.

Our paper has some limitations. Even though we included a wide range of cognitive performance measure in our study, it remains to be tested whether self-reported sleep quality is associated with performance in other cognitive tests, such as attentional or other executive function tasks. It is also possible (as mentioned above), that investigating populations more susceptible to sleep disturbances could yield different results, and the lack of associations could be specific to healthy young adults. Furthermore, it could also be tested if individual differences in other factors (for example, interoceptive ability, i.e., how accurately one perceives their own body sensations) influence the relationship between subjective sleep quality and cognitive performance.

## Conclusions

In conclusion, we showed that self-reported sleep quality is not associated with various aspects of procedural learning, working memory, and executive function in a relatively large sample of healthy young adults. These findings were supported not only by classical (frequentist) statistical analyses, but also by Bayes factors (that provided *evidence for no associations* between these functions). Importantly, however, our findings do not imply that sleep *per se* has no relationship with these cognitive functions; instead, it emphasizes the dissociation between self-reported and objective sleep quality. Together with previous research on dissociations between subjective and objective sleep quality, here we outlined various situations where subjective sleep questionnaires may provide valuable information besides or instead of assessing objective sleep parameters. Nevertheless, careful consideration should be taken in all cases in order to select the best subjective/objective sleep measures depending on the research question. We believe that our approach of systematically testing the relationship between self-reported sleep questionnaires and a relatively wide range of cognitive functions can inspire future systematic studies on the relationship between subjective/objective sleep parameters and cognition.

## Data availability

The datasets analysed during the current study are available in the Open Science Framework repository, https://osf.io/hcnsx/.

## Acknowledgements

This research was supported by the Research and Technology Innovation Fund, Hungarian Brain Research Program (National Brain Research Program, project 2017-1.2.1-NKP-2017-00002); Hungarian Scientific Research Fund (NKFIH-OTKA PD 124148, PI: KJ; NKFIH-OTKA K 128016, to DN); and Janos Bolyai Research Fellowship of the Hungarian Academy of Sciences (to KJ). Authors are thankful to Csenge Török, Kata Horváth, Eszter Tóth-Fáber, Orsolya Pesthy, Noémi Éltető, Andrea Kóbor, and Ádám Takács for their help in data collection.

## Author contributions

Z.Z., J.K. and N.D. designed the present study and wrote the manuscript. G.A. and Z.Z. collected the data. G.A., Z.Z., J.K. and T.N. analyzed the data. Z.Z., J.K., T.N. and N.D. contributed to the interpretation of data and helped revising the previous version of the manuscript critically for important intellectual content. All authors read and approved the final manuscript.

