## Supplementary material for "The relationship between subjective sleep quality and cognitive performance in healthy young adults: Evidence from three empirical studies"

**Supplementary methods**

**Cognitive performance assessments**

***Procedural learning*.** Procedural learning performance was measured by the explicit version of the Alternating Serial Reaction Time (ASRT) task (Figure S1, see also ^1^). In the explicit version of the Alternating Serial Reaction Time (ASRT) task, a stimulus (a dog's head, or a penguin) appeared in one of four horizontally arranged empty circles on the screen, and participants had to press the corresponding button of a Chronos response box (Psychology Software Tools, INC) in Study 1 and Study 2, and a special keyboard with four heightened keys (Z, C, B, and M on a QWERTY keyboard) in Study 3 when the stimulus occurred. The appearance of stimuli followed a predetermined alternating sequence order, such that every second element was part of the sequence and every second element was randomly selected: the dog stimulus always corresponded to sequence elements, and the penguin stimulus indicated random elements (Figure S1A). Participants were informed about this underlying structure of the sequence, and their attention was drawn to the alternation of sequence and random elements by the different visual cues (i.e., dogs vs. penguins). Participants were instructed to respond as quickly and accurately as they could, and also to find the hidden pattern defined by the dog in order to improve their performance.

The task was presented in blocks with 85 stimuli. A block started with five random stimuli for practice purposes, followed by an 8-element alternating sequence that was repeated ten times. The alternating sequence was composed of fixed sequence (pattern) and random elements (e.g., 2-R-4-R-3-R-1-R, where each number represents one of the four circles on the screen and “R” represents a randomly selected circle out of the four possible ones). The timing of the stimulus differed in the three studies. In Study 1, the stimulus remained on the screen until the participant pressed the correct response button, and the next stimulus was presented 250 ms following the previous response. In Study 2, the stimulus remained on the screen until the participant pressed the correct response button, and the next stimulus was presented 120 ms following the previous response. In Study 3, the stimulus remained on the screen for 580 ms, the participant was asked to respond within this time window, and the next stimulus was presented 120 ms following the previous stimulus. Thus, the task was self-paced with different response-to-stimulus intervals (RSI) in Study 1 and 2, while it was fix-paced (inter-stimulus interval, ISI, of 700 ms) in Study 3. The timing parameters of Study 3 was determined based on previous ASRT studies showing that healthy young adults' average RT performance is around 370-430 ms during the task ^1^^,^^2^. We used different settings to explore which timing parameters promote better learning performance. It has been suggested that longer RSI/ISI (e.g., 250 as opposed to 120 ms) can lead to better learning performance as participants have more time to process and elaborate the stimuli ^3^. Nevertheless, it is also plausible that the shorter the time between subsequent stimuli, the easier to find the association among them, which is essential in the ASRT task to achieve a good learning performance.

In all three studies, the ASRT task consisted of 20 blocks. As one block took approximately 1-1.5 min, the session took approximately 20-25 min. For each participant, one of the six unique permutations of the four possible stimulus positions was selected in a pseudo-random manner, so that the six different sequences were used equally often across participants ^4^^,^^5^.

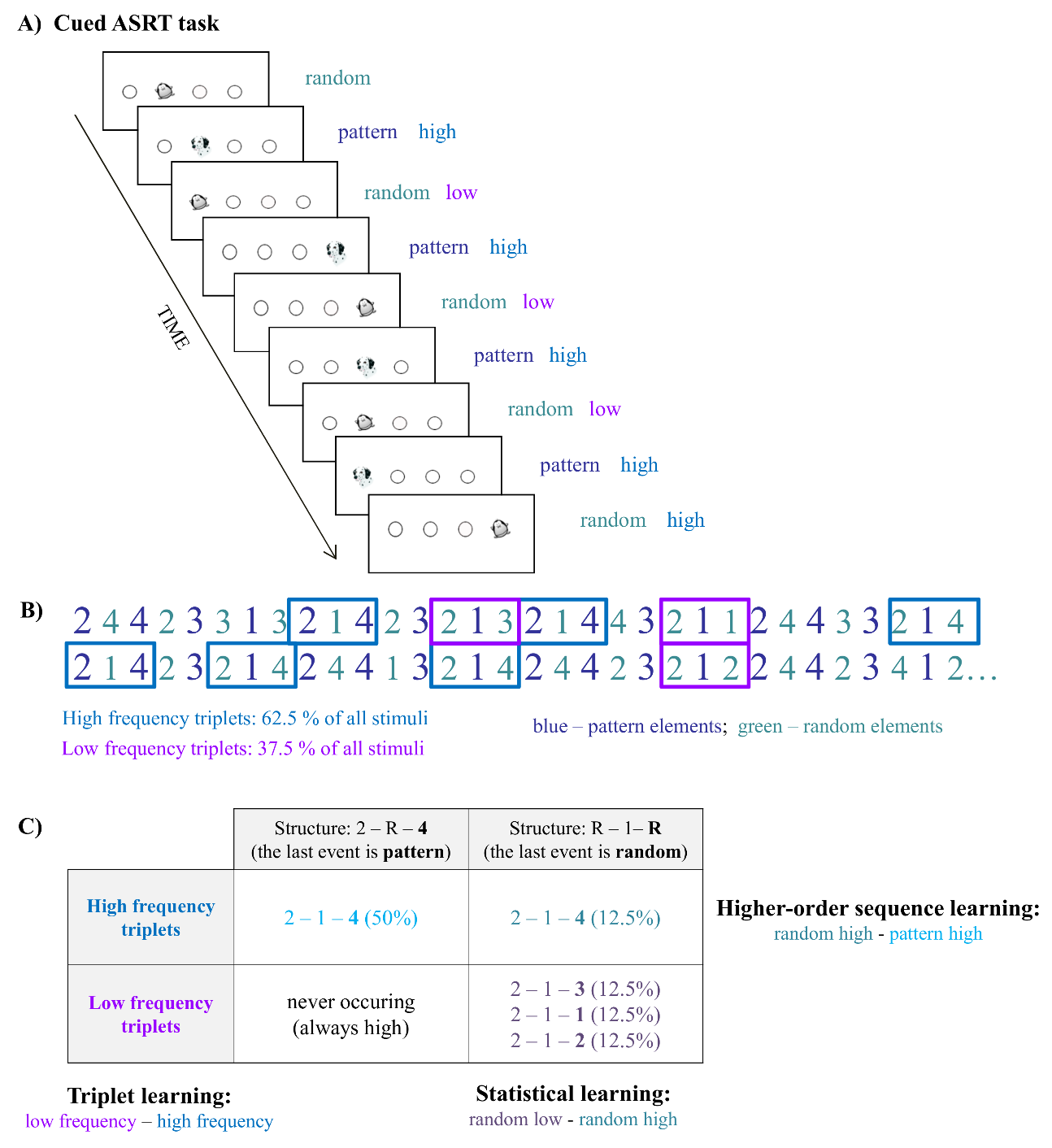

**Figure S1***.* **Schematic diagram of the procedural learning (ASRT) task design**. A) In this task, the appearance of stimuli is based on a predetermined sequence order, in which pattern and random elements alternate (e.g., 2r4r3r1r, where numbers correspond to the four locations on the screen and the 'r' represents randomly chosen locations). The pattern and random elements are cued differently: the dog stimulus always corresponded to pattern elements, and the penguin stimulus indicated random elements. B) Numbers (corresponding to the locations on the screen) in blue represent elements of the pattern trials (e.g., appearing in the sequential order 2, 4, 3, 1 throughout the task), which were alternating with random elements (green). Because of this alternating structure, some runs of three consecutive trials (triplets) occur more frequently than others (high- vs. low-frequency triplets). For each element, we determined whether it was the last element of a high-frequency triplet (one example in blue frame) or low-frequency triplet (examples in magenta frames). C) *Triplet learning* (see text) was calculated as the difference in responses for the last elements of high-frequency triplets (irrespective of random or pattern position) compared to the last elements of low-frequency triplets. *Statistical learning* was assessed by comparing the responses for those *random* elements that were the last elements of high-frequency triplets vs. those that were the last elements of low-frequency triplets (right column). *Higher-order sequence learning* was assessed as a difference between responses for pattern elements (which are always high-frequency triplets) vs. random-high frequency triplet elements (top row).

*Trial types and procedural learning indices in the ASRT task.* The alternating sequence of the ASRT task forms a sequence structure in which some of the runs of three successive trials (henceforth referred to as triplets) appear more frequently than others. In the above example, triplets such as 2X4, 4X3, 3X1, and 1X2 (X indicates the middle element of the triplet) occur frequently since the first and the third elements can either be a pattern or a random stimulus. However, 3X2 and 4X2 occur less frequently since the first and the third elements can only be a random stimulus. Figure S1B and S1C illustrate this phenomenon with the triplet 2-1-4 occurring more often than other triplets such as 2-1-3, 2-1-1, and 2-1-2. The former triplet types are termed as *high-frequency* triplets, whereas the latter types are termed as *low-frequency* triplets (Figure S1C, see also ^1^). The third element of a high-frequency triplet is highly predictable (with 62.5% probability) based on the first element of the triplet. In contrast, in low-frequency triplets, the predictability of the third element is less predictable (with 12.5 % probability) based on the first element of the triplet. According to this principle, each trial was categorized as either the third element of a high- or a low-frequency triplet.

Additionally, trials are differentiated by the visual cues (dog vs. penguin) indicating whether a pattern or a random stimulus was presented in that given trial. In case of pattern trials, participants can use their explicit knowledge of the sequence to predict that trial. Consequently, we further differentiate the previously defined high-frequency triplets into two categories based on whether the last element of the triplet was a pattern or a random stimulus. This way, the task consists of three trial types: 1) trials that belong to the explicitly cued sequential pattern and, at the same time, appear as the last element of a high-frequency triplet are termed *pattern* trials; 2) trials of random stimuli that appear as the last element of a high-frequency triplet are termed *random high* trials; and 3) trials of random stimuli that appear as the last element of a low-frequency triplet are termed *random low* trials (see the example in Figure S1C).

Previous studies have shown that as people practice the ASRT task, they come to respond more quickly and more accurately to the high-frequency triplets (irrespective of whether it was for a pattern or a random stimulus) compared to low-frequency triplets (always random), revealing *Triplet learning* ^5^^,^^6^. *Triplet learning* is measured as the difference in reaction time (RT) and accuracy (ACC) between high- and low-frequency triplets (RTs of low-frequency triplets minus RTs of high-frequency triplets; ACC of high-frequency triplets minus ACC of low-frequency triplets). Thus, greater Triplet learning is defined as faster/more accurate responses to high-frequency triplets compared to low-frequency triplets. Importantly, however, the comparison of RT and ACC of high- vs. low-frequency triplets does not take into account whether the last elements of the high-frequency triplets are pattern or random stimuli and consequently, provides a mixed measure of at least two separate learning processes.

The two key learning processes that can be disentangled in the explicit ASRT task are the so-called *Higher-order sequence learning* and the so-called *Statistical learning* (Figure S1C). *Higher-order sequence learning* is measured as the difference in RTs between random high and pattern trials (RTs for random high trials minus RTs for pattern trials; ACC for pattern trials minus ACC for random high trials). These trials share the same statistical properties (both correspond to the third element of high-frequency triplets) but have different sequence properties (i.e., pattern vs. random trials). Thus, greater Higher-order sequence learning is defined as faster/more accurate responses to pattern trials compared to random high trials. This learning measure thus can reflect the knowledge about the alternating sequential structure that the participants explicitly acquired during the task.

*Statistical learning* is assessed by comparing the responses for those random trials that were the last elements of a high-frequency triplet vs. those that were the last elements of a low-frequency triplet (RTs for random low trials minus RTs for random high trials; ACC for random high trials minus ACC for random low trials). These trials share the same sequence properties (both are random) but differ in statistical properties (i.e., they correspond to the third element of a high- or a low-frequency triplet). Hence, faster responses to random high compared to random low trials yields greater Statistical learning. While Higher-order sequence learning quantifies the acquisition of the sequential pattern, Statistical learning captures purely frequency-based learning ^1^^,^^7^. Based on previous findings, the cueing of pattern and random stimuli is necessary to promote Higher-order sequence learning, otherwise, it occurs more slowly, and cannot be acquired during a single session ^1^^,^^4^.

Additionally, more general changes in RT and ACC performance can be measured in the ASRT task. These changes occur similarly for all trial types, thus are not related to acquiring the sequential or statistical structure embedded in the stimulus stream. Instead, these general changes indicate general skill improvements, such as more efficient visuo-motor and motor-motor coordination as the task progresses ^8^, combined with potential fatigue effects that can accumulate during practice ^9^^,^^10^. General skill improvements in terms of RT are assessed as the difference of speed in the beginning and at the end of the task (RTs of the first five blocks minus RTs of the last five blocks, see also Statistical analysis). Similarly, general changes can be quantified in ACC between the beginning and the end of the task (ACC of the first five blocks minus ACC of the last five blocks).

In our study, we first report the Triplet learning results because this has been the most common analysis method in the ASRT studies and thus it enables to directly compare our results with those of previous studies. Next, we report Higher-order sequence learning and Statistical learning measures to obtain a more detailed picture of the underlying processes within procedural learning. Finally, we report average RTs and ACCs and their change from the beginning to the end of the end of the task to test whether these more general aspects of performance have a differential association pattern with sleep compared to the learning scores.

***Working memory.*** The Counting Span task ^11-14^ was used to assess working memory (WM) performance. The task consisted of three series. In each series, each trial included three to nine blue circles as targets, one to nine blue squares, and one to ﬁve yellow circles as distractors on a grey background. Participants counted aloud the number of blue circles in each trial, and when ﬁnished with counting, they repeated the total number. When presented with a recall cue, participants recalled each total from the preceding set of trials, in the order in which they appeared. The number of presented trials (i.e., set) ranged from two to six. A participant’s counting span capacity was calculated as the average of the highest set sizes of the three series at which the participant was able to recall the totals in the correct serial order.

***Executive functions.*** The Wisconsin Card Sorting Test (WCST, ^15^^,^^16^) was used to assess executive functions. In this task, participants are asked to find out a sorting rule for cards based on the feedback they receive for their card-sorting choices. During the task, there are four decks on the screen with symbols on them, which differ in three features: number, shape, and color. On the bottom of the screen, a stimulus card appears, and participants have to match this card to one of the decks (based on a sorting rule of their choice). After the choice, participants receive a feedback whether the choice was correct or not. Based on the feedback, participants have to find out the correct sorting rule. In each trial, only one sorting rule is correct (e.g., number, shape or color), and the rule changes several times during the task, allowing to measure adaptation to changing rules. The outcome measure of the task is the number of *perseverative errors*, which shows the inability to change the behavior despite feedback, so the higher values of this measure indicate weaker executive functions.

**Subjective sleep quality assessments**

***Athens Insomnia Scale.*** The Athens Insomnia Scale (AIS, ^17^^,^^18^) was administered in all three studies. AIS is a self-reported questionnaire assessing general sleep quality (over a one month time period), and consists of eight items; the first five items assess difficulty with falling asleep, awakening during the night, early morning awakening, total sleep time, and overall quality of sleep, while the last 3 items pertain to the sense of well-being, overall functioning and sleepiness during the day. Each item of AIS can be rated from 0 to 3 and the total score ranges from 0 to 24, where higher scores indicate poorer sleep quality.

***Pittsburgh Sleep Quality Index.*** The Pittsburgh Sleep Quality Index (PSQI, ^19^^,^^20^) is one of the most commonly used questionnaires measuring self-reported sleep habits and sleep disturbances over the last month. Here we focused on three components of the questionnaire, which were obtained in all three studies: subjective sleep quality, sleep latency and sleep disturbances. Item 6 (referring to the original coding of PSQI) measured the participant’s perceived sleep quality, item 5a indicated sleep latency and items 5b-5j (9 items) showed sleep disturbances. We chose the aforementioned items because the factors they form are those that contribute most to the overall PSQI score (besides daytime dysfunction which we didn’t include, because we wanted to measure daytime functioning with the tests included). These three components range from 0 to 3 and form a global score that ranges between 0 and 9, a higher score indicating poorer sleep quality. Henceforth we refer to this 11-item long PSQI as PSQI.

***Groningen Sleep Quality Scale.*** In Study 2, subjective sleep quality of the night before cognitive testing was assessed by the Groningen Sleep Quality Scale (GSQS, ^21^^,^^22^), which is a 15-item self-administered questionnaire. Every item is a yes or no question, scoring 0 or 1, thus GSQS scores range from 0 to 14 (the first item is typically not scored), a higher score indicating poorer quality of sleep.

***Sleep diary.*** In Study 2, we also asked participants to keep a sleep diary for 1-2 weeks prior the testing session ^23^. In this diary, participants had to mark the time they went to bed, the time they got up, and the hours they spent with sleep during this period. After each night, participants also had to rate how good their sleep was (on a scale from 1 to 5), report how long it took them to fall asleep (in minutes), and how many times they woke up during the night. We evaluated data from sleep diaries similarly to PSQI component scores. The average subjective sleep quality was scored as the first component of PSQI; the average sleep latency as the second component of PSQI, the average time spent with sleep as the third component of PSQI, and the average sleep time divided with the time spent in bed (i.e., sleep efficiency) as the fourth component of PSQI. Altogether, based on the sleep diary, we had 4 component scores, ranging from 0 to 3 and form a global score that ranges between 0 and 12, a higher score indicating poorer quality of sleep.

**Statistical analysis**

***Analysis of the ASRT data.*** To facilitate data processing and to reduce intra-individual variability, the blocks of ASRT were collapsed into epochs of five blocks, following previous ASRT studies ^1^^,^^14^. The first epoch contained blocks 1–5, the second epoch contained blocks 6–10, etc. We calculated mean accuracy (ACC) for all responses, and median reaction times (RTs) for correct responses only, separately for pattern, random high and random low trials for each epoch. As in previous ASRT studies ^5^, two kinds of low-frequency triplets were eliminated: repetitions (e.g. 222, 333) and trills (e.g. 212, 343). Repetitions and trills are low-frequency for all participants, and people often show pre-existing response tendencies to them ^24^. By eliminating these triplets, we attempted to ensure that differences between high- vs. low-frequency triplets emerged due to learning and not to pre-existing response tendencies.

Performance in the ASRT task was analyzed by repeated measures analyses of variance (ANOVA) with median RTs or mean ACCs as the outcome measure, and EPOCH (1^st^-4^th^ epochs) and TRIAL TYPE (pattern, random-high, and random-low) as within-subject factors. To evaluate the effect of TRIAL TYPE, and thus to confirm that Higher-order sequence learning (pattern vs. random-high difference) and Statistical learning (random-high vs. random-low difference) occurred, Fisher’s LSD post-hoc comparisons were performed. Greenhouse-Geisser epsilon (ε) correction was used if necessary. Original *df* values and corrected *p* values (if applicable) are reported together with partial eta-squared (η*_p_*^2^) as a measure of effect size. Note that for the sake of brevity, we do not report ANOVAs for the mixed measure of Triplet learning (RT/ACC difference in responses to high- vs. low-frequency triplets), which does not clearly differentiate between acquiring the sequential and statistical structure embedded in the task. We conducted these ANOVAs, and Triplet learning occurred in all three studies, both in ACC and RT (significant main effect of TRIPLET: all *p*s < .001).

**Supplementary results**

**Working memory and executive function performance in the three studies**

The working memory capacity, and executive functions (measured as number of perseverative errors in the WCST task) of participants were in the standard range for their age ^25^^,^^26^. The mean counting span score for the entire sample was 3.59 (SD = 0.85) in the three studies. These average scores represent a mid-range cognitive performance, as the maximum obtainable score is 6. Mean of the number of perseverative errors was 14.76 (SD = 5.27) in the three studies (no maximum score can be defined in this case). For procedural learning, mean scores were 26.48 (SD = 26.37) for RT Triplet learning, 16.63 (SD = 40.34) for RT Higher-order sequence learning, 16.74 (SD = 9.94) for RT Statistical learning, 359.88 (SD = 40.94) for average RT, and 31.13 (SD = 30.15) for RT general skill learning. Accuracy scores were as follows: 0.04 (SD = 0.03) for ACC Triplet learning, 0.02 (SD = 0.03) for ACC Higher-order sequence learning, 0.03 (SD = 0.03) for ACC Statistical learning, 0.90 (SD = .10) for average ACC, -0.02 (SD = 0.09) for ACC general skill learning, in all three studies. Note that in case of accuracy, these values represent proportions (e.g., the average ACC was 90%, hence 0.90), and the learning scores are difference score (e.g., the ACC Triplet learning score shows that participants were on average 4% more accurate on high-frequency triplets compared to the low-frequency ones). All presented RT and ACC scores represent typical values in ASRT studies with healthy young adults.

We also provide descriptive data for Study 2 separately, as additional analyses were run on cognitive performance from this dataset and GSQS and sleep diary scores. In Study 2, mean counting span score was 3.65 (SD = 1.01), and mean of the number of perseverative errors was 14.46 (SD = 6.37). For procedural learning in Study 2, mean scores were 33.04 (SD = 27.96) for RT Triplet learning, 28.53 (SD = 51.44) for RT Higher-order sequence learning, 18.77 (SD = 9.78) for RT Statistical learning, 348.29 (SD = 42.26) for average RT, and 39.30 (SD = 34.74) for RT general skill learning. Accuracy scores were as follows: 0.03 (SD = 0.02) for ACC Triplet learning, 0.01 (SD = 0.02) for ACC Higher-order sequence learning, 0.02 (SD = 0.02) for ACC Statistical learning, 0.94 (SD = 0.03) for average ACC, 0.02 (SD = 0.03) for ACC general skill learning. These standard deviations show reasonable variability in the sample to conduct the planned analyses.

**Subjective sleep questionnaire scores in the three studies**

Mean AIS score in the three studies were 3.98 (SD = 2.66) and 3.41 (SD = 2.09) in Study 2. Overall, these average scores constitute the lower portion of the AIS scale, as scores can range from 0 to 24, with higher scores indicating poorer sleep quality. Mean PSQI scores were 2.99 (SD = 1.57) in all three studies, and 2.54 (SD = 1.29) in Study 2. As the obtainable scores on PSQI range between 0 and 9, these average scores represent the lower half of the PSQI scale, although proportionally are somewhat higher (and thus suggest a slightly poorer sleep quality) compared to the AIS scores. The aggregated sleep disturbance index (that incorporates the AIS and PSQI scores) over all studies ranged from -1.9 to 3.86. Mean of the GSQS score in Study 2 was 2.86 (SD = 2.87) on the scale of 0 to 14. Mean of the Sleep diary score in Study 2 was 1.38 (SD = 1.22) on the scale of 0 to 12. Altogether, these scores suggest a relatively good sleep quality in the healthy young adult populations of Study 1-3. These standard deviations show reasonable variability in the sample to conduct the planned analyses.

**Procedural learning across the three studies**

***Study 1.*** The repeated-measures ANOVA on RT data (Figure S2A) revealed a significant main effect of EPOCH (*F*_3,138_ = 33.84, *p* < .0001, η_p_^2^ = .42), such that RTs decreased as the learning progressed indicating general skill improvements. The main effect of TRIAL TYPE was significant as well (*F*_2,92_ = 60.89, *p* < .001, η_p_^2^ = .57). The post-hoc analysis revealed that responses (averaged across epochs) to pattern trials were faster (M = 340.27 ms) compared to random high (M = 348.70 ms, *p* = .002) and random low trials (M = 366.68 ms, *p* < .0001), and responses to random high trials were faster compared to random low trials (*p* < .0001). The different RTs for the different trial types indicate Higher-order sequence learning (difference in pattern vs. random high trials) and Statistical learning (difference in random low vs. random high trials) occurred during the task. The interaction between EPOCH and TRIAL TYPE was not significant (*F*_6,276_ = .89, *p* = .41, η_p_^2^ = .02).

The repeated-measures ANOVA on accuracy data (Figure S2B) again revealed a significant main effect of EPOCH (*F*_3,138_ = 13.08, *p* < .0001, η_p_^2^ = .22): ACC decreased during the task. Similarly, to RTs, there was a significant main effect of TRIAL TYPE (*F*_2,92_ = 51.06, *p* < .0001, η_p_^2^ = .53). The post-hoc analysis revealed that responses (averaged across epochs) to pattern trials were more accurate (M = 96%) compared to random high (M = 95%, *p* = .08) and random low trials (M = 93%, *p* < .0001), and responses to random high trials were more accurate compared to random low trials (*p* < .0001). This again indicates that Higher-order sequence learning and Statistical learning occurred during the task. The EPOCH x TRIAL TYPE interaction was also significant (*F*_6,276_ = 4.15, *p* = .001, η_p_^2^ = .08), the ACC for random low trial type decreased more during the task (3.4% decrease on average) than ACC for pattern (1.2% decrease) or random high trial types (1% decrease).

***Study 2.*** Similarly to Study 1, the ANOVA for RTs (Figure S2C) revealed a significant main effect of EPOCH (*F*_3,306_ = 93.13, *p* < .0001, η_p_^2^ = .48): RTs decreased as the learning progressed. We also found a significant main effect of TRIAL TYPE (*F*_2,204_ = 63.39, *p* < .0001, η_p_^2^ = .38). The post-hoc analysis revealed that responses (averaged across epochs) to pattern trials were faster (M = 323.01 ms) compared to random high (M = 351.55 ms, *p* = .002) and random low trials (M = 370.32, *p* < .001), and responses to random high trials were faster compared to random low trials (*p* < .001), indicating Higher-order sequence learning and Statistical learning occurred during the task. The EPOCH x TRIAL TYPE interaction was also significant (*F*_6,612_ = 8.74, *p* < .001, η_p_^2^ = .08), RTs for pattern trials decreased more (58 ms on average) than for random high (31 ms) and random low (28 ms) trials.

Again, in the case of accuracy (Figure S2D), there was a significant main effect of EPOCH (*F*_3,306_ = 18.12, *p* < .0001, η_p_^2^ = .15): ACC decreased during the task. There was also a significant main effect of TRIAL TYPE (*F*_2,204_ = 97.55, *p* < .0001, η_p_^2^ = .49). The post-hoc analysis revealed that responses (averaged across epochs) to pattern trials were more accurate (M = 96%) compared to random high (M = 95%, *p* < .001) and random low trials (M = 92%, *p* < .001), and responses to random high trials were more accurate compared to random low trials (*p* < .001), indicating Higher-order sequence learning and Statistical learning occurred during the task. The EPOCH x TRIAL TYPE interaction was also significant (*F*_6,612_ = 5.47, *p* = .0001, η_p_^2^ = .05), the ACC for random low trial type decreased more during the task (3.3%) than ACC for pattern (0.7% decrease) or random high trial (1.3% decrease) types.

***Study 3.*** Similar to the previous studies, in the case of RT data (Figure S2E), there was a significant main effect of EPOCH (*F*_3,252_ = 45.60, *p* < .0001, η_p_^2^ = .35): RTs decreased during the task. The main effect of TRIAL TYPE was significant as well (*F*_2,168_ = 31.58, *p* < .0001, η_p_^2^ = .27). The post-hoc analysis revealed that responses (averaged across epochs) to pattern trials were faster (M = 367.03 ms) compared to random high (M = 375.92 ms, *p* = .008) and random low trials (M = 388.93, *p* < .0001), and responses to random high trials were faster compared to random low trials (*p* < .0001). The different RTs for the different trial types again indicate that Higher-order sequence learning and Statistical learning occurred during the task. The EPOCH x TRIAL TYPE interaction was also significant (*F*_6,504_ = .4.50, *p* < .001, η_p_^2^ = .05), the RTs for pattern and random high trial types decreased more (27 ms and 22 ms respectively) than RTs for random low trials (13 ms).

Again, the ANOVA on accuracy data (Figure S2F) revealed a significant main effect of EPOCH (*F*_3,252_ = 33.03, *p* < .0001, η_p_^2^ = .28), such as ACC decreased during the task, and a significant main effect of TRIAL TYPE (*F*_2,168_ = 78.97, *p* < .001, η_p_^2^ =.49). The post-hoc analysis revealed that responses (averaged across epochs) to pattern trials were more accurate (M = 84%) compared to random high (M = 82%, *p* < .0001) and random low trials (M = 78%, *p* < .0001), and responses to random high trials were more accurate compared to random low trials (*p* < .0001) indicating that Higher-order sequence learning and Statistical learning occurred during the task. The EPOCH x TRIAL TYPE interaction was not significant (*F*_6,504_ = .91, *p* = .47, η_p_^2^ = .01).

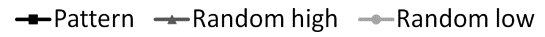

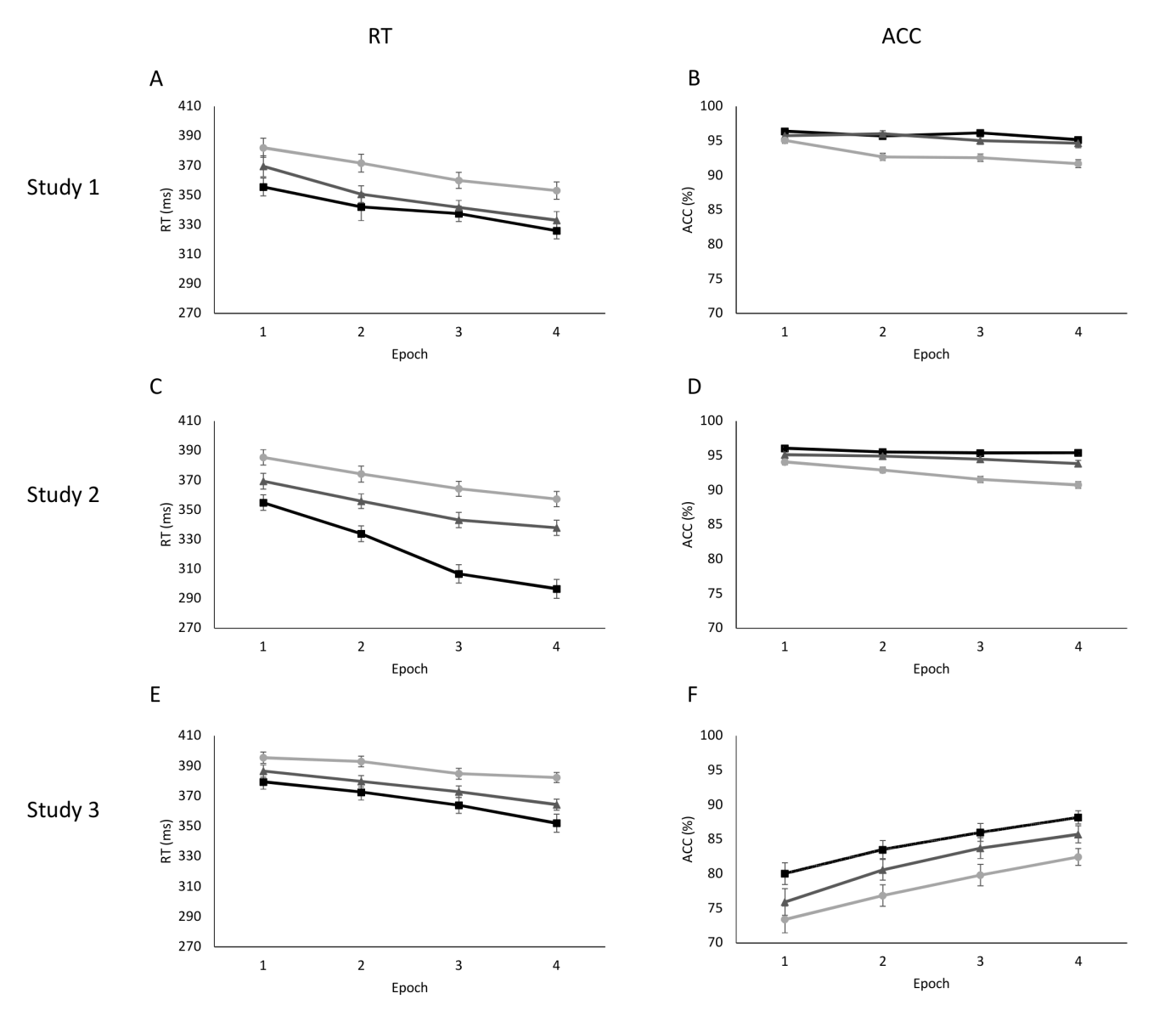

**Figure S2.** RT for correct responses (A, C, E) and accuracy for all responses (B, D, F) as a function of epoch (1-4) and trial type (pattern, random high- and low-frequency trials) in the ASRT task assessing procedural learning. The gap between the curves of pattern and random high-frequency trials indicates Higher-order sequence learning, the gap between the curves of random high and low-frequency indicates Statistical learning. Error bars denote standard error of mean.
